# Assessing the effect of zoo closure on the soundscape using multiple measures

**DOI:** 10.1101/2023.05.19.540934

**Authors:** Rebecca N Lewis, Leah J Williams, Selvino R de Kort, R Tucker Gilman

## Abstract

The zoo soundscape has a number of important implications for animal welfare, management, and conservation. However, despite its importance, the zoo soundscape is yet to be examined in depth using multiple measures. Consistent human presence can influence the zoo soundscape. However, it is difficult to determine the specific impact of human presence, as visitors are usually present during the day when animals are active. The COVID-19 lockdown in 2020 provided a unique opportunity to study zoo soundscapes in the absence of visitors. We compared the sound environment during the 2020 closure period to a comparable open period in 2019 across three zoo aviaries, examining broad band frequency measures of sound pressure levels, sound pressure levels in defined frequency bands, and acoustic indices (Acoustic Complexity Index and Normalized Difference Soundscape Index) to describe the zoo soundscape. Acoustic indices have not, to our knowledge, previously been used in the zoo setting, although they may provide a useful metric to assess sound disturbance. Therefore, we also used this natural experiment to explore how successful these measures may be in assessing disturbances in captive environments. We found a significant effect of human presence on the sound environment; aviaries were generally quieter with less low frequency noise and with a greater proportion of biotic sound during the 2020 zoo closure period. We argue that NDSI could be a useful index for determining anthropogenic disturbance in zoos, although further information on how it is influenced by additional factors, such as human speech, would be beneficial. The use of multiple measures to assess the sound environment in zoos can provide additional information beyond ‘loudness’, such as frequencies where sound energy is concentrated and characteristics of the soundscape, which could be used to better target management and mitigation.

## 1. Introduction

Zoos differ from wild environments in a number of aspects, such as the lack of predators, reductions in disease and parasites, and space constraints (Frankham 2008). Perhaps the clearest difference between zoo environments and those in the wild is the consistent presence of human influences. The presence of human visitors, often measured by the overall number of visitors, on animal behaviour and welfare has been widely examined (Hosey 2000, 2008; Davey 2007; Fernandez et al. 2009; Sherwen & Hemsworth 2019). However, the effect of human presence on other aspects of the environment, such as environmental sound, is less often considered.

Human presence can influence the soundscape experienced by animals in zoos. In general, sound sources in zoos can be separated into four broad categories: 1) permanent sources of anthropogenic sound, such as heating, ventilation, and air conditioning (HVAC) systems; 2) temporary anthropogenic sound from maintenance, construction, and other events; 3) human speech and footfall; and 4) sounds produced by other animals (Clark & Dunn 2022). Three of these four categories are related to human presence and anthropogenic sound sources, suggesting that the constant presence of humans is likely to have a significant impact on the soundscape. Sound produced by other animals may also vary between wild and captive environments due to novel species composition in zoos. As such, animals in captivity may experience soundscapes that differ from those in the wild (Lara & Vasconcelos 2019). In addition, the contributions of different sound sources are likely to differ among zoos and enclosures, and, therefore, soundscapes are also likely to vary among captive environments (Clark & Dunn 2022).

The soundscape is an important consideration in captivity due to its potential effects on animal behaviour, communication, and welfare. Increases in sound from visitors have been associated with a wide range of behavioural changes across taxa, which are often interpreted as negative (reviewed in Sherwen & Hemsworth, 2019). As well as potential effects on animal behaviour, exposure to environmental sound can also affect physiology, development, neural functions, and genetics (Kight & Swaddle 2011), and may even lead to hearing impairment (Wolfenden et al. 2019). Importantly for ex situ breeding programmes, sound can also negatively affect reproductive success (Halfwerk et al. 2011; Kight et al. 2012). Increased background sound can obscure sounds that are important for survival and reproduction, an effect known as masking, which could reduce communication efficacy (Brumm & Slabbekoorn 2005; Barber et al. 2010; Blickley & Patricelli 2010). Many taxa are reported to alter vocal behaviour to reduce the effect of masking (Brumm & Slabbekoorn 2005; Slabbekoorn & Ripmeester 2008). As such, the soundscape in captivity may alter vocal behaviour, contributing to vocal divergence between populations, which may have knock-on effects for conservation programmes (Passos et al. 2017; Lewis et al. 2021). Given its potential effects on animal behaviour, welfare and conservation, understanding the soundscape has important implications for animal management in captivity.

Despite its importance, the study of sound in zoos has lagged behind that in wild environments (Clark & Dunn 2022) and sound is less often considered than other features of the zoo environment (Binding et al. 2020). Past work has focused mostly on measuring maximum sound pressure levels, which provide a quantification of environmental sound in decibels, most often in dB(A) (Clark & Dunn 2022). This metric is only concerned with the level of environmental sound within the human hearing range, but does not provide any additional information about the spectral characteristics of the soundscape overall i.e., what that sound is like. However, there is increasing interest in sound in zoological collections (Pelletier et al. 2020; Rose et al. 2021; Clark & Dunn 2022). Due to recent technological advances, such as the development of low-cost autonomous recording units (ARUs), it is possible to collect large amounts of data with comparatively little effort (Brandes 2008). This allows for a more in-depth study of the soundscape beyond traditional sound levels, such as the investigation of sound in specific frequency bands (Pelletier et al. 2020). There also is ample opportunity to use measures such as acoustic indices, which were developed for use in the wild, in captivity (Sueur et al. 2014; Bradfer-Lawrence et al. 2019; Clark & Dunn 2022). Acoustic indices reduce multidimensional data into a single metric which provides information about the characteristics of sound in the environment, such as complexity or diversity, rather than just measuring sound intensity (Sueur et al. 2014). Although sound pressure level is an important measure for examining the soundscape and its effects on animals in zoos, studying additional measures such as acoustic indices can give a more well-rounded view of sound in zoos, which will be beneficial for management.

As zoos are rarely closed for prolonged periods, visitors are present throughout the year. Therefore, it is difficult to determine the effects of human presence on the soundscape, as humans are constantly present during the day when many animals are active. The COVID-19 lockdown in 2020 provided a unique opportunity to study sound in the zoo in the absence of visitors and with reduced influences from other anthropogenic sound sources e.g., road and air traffic. A number of studies have examined the effect of lockdown and the absence of visitors on animal behaviour in zoos (Carter et al., 2021; Finch et al., 2022; Jones et al., 2021; Masman et al., 2022; Podturkin, 2022; Williams et al., 2021b, 2021a, 2022), but, to our knowledge, the effect on the soundscape has not been explored. We examined the soundscape in three mixed species aviaries to explore the effect of human presence. We examined sound pressure levels using broad frequency band measures (dB(A) and 10 Hz – 10 kHz dB(Z)), as well as in narrower, defined frequency bands (dB(Z)). In addition to sound pressure levels, we also examined two acoustic indices; the Acoustic Complexity Index (ACI) (Pieretti et al. 2011), which measures soundscape complexity, and the Normalized Difference Soundscape Index (NDSI) (Kasten et al. 2012), which quantifies soundscape naturalness. As acoustic indices have not been used in zoos, this natural experiment also provides the opportunity to examine the effectiveness of these indices in monitoring disturbance in captivity.

## 2. Methods

### 2.1 Data collection sites and periods

Data were collected from three aviaries at Chester Zoo, UK, between April 30^th^ and May 21^st^ 2019 and April 8^th^ and May 9^th^ 2020 (see Supplementary Information 1 for precise dates). The Bali Temple aviary is an outdoor, walkthrough aviary located near the edge of the zoo, in an open area close to a main road (A41). The Sumatra aviary is also an outdoor, walkthrough aviary, near to the Bali Temple, but in a sheltered location surrounded by zoo buildings. In a walkthrough aviary, the visitor path is located inside the aviary, i.e., visitors and birds occupy the same space without barriers. The Dragons in Danger aviary is an indoor aviary located at the centre of the zoo. Although this aviary is not a walkthrough, the visitor path moves through the building with netted aviaries either side of the walkway, meaning that there is no sound barrier between visitors and birds. Each aviary contained a range of bird species, including various passerines, pigeons, pheasants and parrots, many of which are native to South East Asia (species lists for each aviary in each time period can be found in Supplementary Information 1).

### 2.2 Data collection method

Sound recordings were made using Wildlife Acoustics SM4s (Wildlife Acoustics Inc., Maynard, MA, USA) placed within the enclosure. For the Dragons in Danger aviary, the recording device was placed in a central location within the bird area, although this was outside of the netted aviaries. Devices were set to record continuously across the days, and continued to record until either the batteries ran out or the memory card reached capacity. We used a 24 kHz sampling rate for recordings. This allowed us to capture information on sounds below 12 kHz, which is appropriate for birds, as biotic sound (and the frequency limits of many bird songs) is typically concentrated below 8 kHz (Slabbekoorn & Ripmeester 2008; Pijanowski et al. 2011; Kasten et al. 2012).

### 2.3 Data extraction

Data for sound pressure levels and acoustic indices were extracted using Kaleidoscope Pro 5.4.7 (Wildlife Acoustics Inc., Maynard, MA, USA).

#### 2.3.1 Broad frequency band measures

To assess sound pressure levels (broad and defined bands), we extracted the mean (L_eq_) sound pressure level (SPL) for each 1-hour period during the day. Mean sound pressure level is a representative measure for sounds that remain more or less constant over time, and is only weakly affected by sharp bursts of sound that may occur e.g., slamming doors. Sound pressure levels used the standard reference of 20uPa, where 1 Pa is equal to a sound pressure level of 94 dB.

We extracted two broad frequency band measurements, A-weighted decibels (dB(A)) across the recording, and Z-weighted decibels (dB(Z)) between 10 – 10000 Hz. A-weighted sound level measurements apply a filter adjusted to the human hearing range with reduced weighting of infra- and ultra sounds (Kurra 2021). A-weighted measurements are often used in acoustic studies in zoos, as most commercial sound level meters use this metric. Therefore, we included dB(A) values to allow for comparison with other studies. Moreover, birds’ hearing range is broadly similar to that of humans (Catchpole & Slater 2008) suggesting that dB(A) may be meaningful when examining sound in an aviary setting. Z-weighted dB uses a flat-frequency response, where all frequencies are given equal weight, providing a less anthropocentric measure of sound in the environment. Therefore, dB(Z) provides a broader measure of sound in the environment, which is not based on the hearing range of a particular species.

#### 2.3.2 Defined frequency band measures

We also examined sound levels in different frequency bands. Understanding how sound energy is distributed in the environment can help us to identify which sounds are prevalent, determine which species are most likely to be affected by noise, and appropriately target mitigations, which often act across a limited range of frequencies (Orban et al. 2017). We extracted information on sound pressure levels from 30 third-octave bands (central frequencies 19.7 – 10079.4 Hz). We then combined the sound pressure level (SPL) from these bands into four larger frequency bands for analysis. Our categories were defined based on their relevance for species and sound in zoos: very low frequency (17.6-111.4 Hz); low frequency (111.4 – 890.9 Hz); mid frequency (890-9 – 8979.7 Hz); and high frequency (8979.7 – 11313.7 Hz). Each third-octave band was given equal weighting to all bands, providing a measure in dB(Z), using the following equation, where the summation runs over all third-octave bands in the bin being created (Lin et al. 2021):

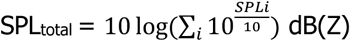

These divisions are meaningful for the study of sound in aviaries and elsewhere in zoological environments. The very low frequency band covers sound below 100 Hz, including some infrasound (<20 Hz), which can effect health in both humans and animals (Persinger 2014; Lousinha et al. 2018; Pereira et al. 2021). Many bird taxa use frequencies below 1 kHz for communication (Slabbekoorn & Ripmeester 2008), and therefore may contribute to and be impacted by sound in the ‘Low’ frequency band. The ‘Mid’ frequency band is biologically relevant for many species, covering a large proportion of biotic sounds (Pijanowski et al. 2011; Kasten et al. 2012) and the sensitive hearing ranges of most mammals (Fay, 1988) and birds (Dooling 1992; Dooling et al. 2000). Some birds may also use higher frequencies in songs and for alarm calls, so will contribute to and be impacted by the ‘High’ band. Our combined frequency bands also align with those calculated in Pelletier et al. (2020), which facilitates comparisons and increases repeatability (Clark & Dunn 2022). However, unlike Pelletier et al. (2020), we only studied sounds below 12 kHz, as these are likely to be most relevant for birds.

#### 2.3.3 Acoustic indices

Acoustic indices are statistics that can be used to summarize aspects of the soundscape (Sueur et al. 2014; Towsey et al. 2014). Acoustic indices have not previously been used in zoo environments (Clark & Dunn 2022), but could be useful in understanding sound in zoos beyond sound pressure levels. We identified two indices, detailed below, that may be particularly relevant to the zoo environment and examined how they behave in the different aviaries and situations in this study. For each index we used the standard settings (Pieretti et al. 2011; Kasten et al. 2012), as outlined in the original papers, during extraction. Whilst acoustic indices can be tuned to match individual habitats, we chose to use the standard measures for two reasons. Firstly, acoustic indices are yet to be used in zoo settings, so it is difficult to determine how default settings perform and, if sub-optimally, how they should be adjusted. Secondly, there is a need for consistent measurements across zoos so that comparisons can be made (Clark & Dunn 2022). The use of default settings means methods are easily replicable between collections. For each index, we extracted index values for each 60 s section of recordings. An average of each index was then taken per hour to be used in analyses.

We extracted the Acoustic Complexity Index (ACI) (Pieretti et al. 2011), which is based on the observation that many biotic sounds are intrinsically variable, whilst anthropogenic sound is often more constant (Pieretti et al. 2011; but see Wolfenden et al. 2019). The index compares amplitude differences between time intervals within narrow frequency bands, and these measures are combined across frequency bands to calculate the overall index (Pieretti et al. 2011). Natural soundscapes, e.g., those with high biophony (sounds of biotic origins e.g., birds), have high ACI values (Pieretti et al. 2011). In wild environments, ACI values have been found to correlate with the number of bird vocalizations (Pieretti et al. 2011). We used the entire frequency range of the recording to calculate the ACI using an FFT window size of 552 (Pieretti et al. 2011). Examining diel patterns in ACI could indicate whether the metric responds appropriately to increases in vocal activity during daylight hours in the zoo. In addition, by comparing ACI values in 2019 and 2020, we can assess whether soundscape complexity is affected predictably by anthropogenic disturbances.

The Normalized Difference Soundscape Index (NDSI) (Kasten et al. 2012) determines the relative ratio of anthropogenic compared to biotic sound, providing a useful metric to assess the dominance of anthropogenic sounds in captive environments. Anthropogenic sounds are generally prevalent between 1 and 2 kHz, although many anthropogenic sounds also occur below 1 kHz, whereas biological sounds are more commonly found between 2 and 8 kHz (Pijanowski et al. 2011; Kasten et al. 2012). Therefore, these bands are used to represent anthropogenic (1 – 2 kHz) and biotic (2 – 8 kHz) sound in the index. Some sounds in our recordings, particularly at low frequencies, may fall outside of this range, but we chose to use the default values for acoustic indices to aid repeatability. Briefly, the NDSI calculates the power spectral density of the signal and estimates of the power spectral density integral are computed for each of the specified frequency ranges, which are then compared using the formula NDSI = (*β* − *α*)/(*β* + *α*), where *β* and *α* are the total estimated power spectral density for the largest 1 kHz biophony and anthrophony bin respectively (Kasten et al. 2012). The NSDI returns a value between −1 and +1, with positive values indicating relatively more biotic compared to anthropogenic sound and negative values indicating relatively more anthropogenic compared to biotic sound (Kasten et al. 2012). We used an FFT window size of 512. By comparing the 2019 and 2020 periods, we can assess how NDSI changes with anthropogenic disturbances and determine its effectiveness as a metric for anthropogenic noise disturbance in zoos.

### 2.4 Data analysis

To examine the effect of visitor absence on features of the soundscape we used a series of Generalized Additive Models (GAMs) implemented in the mgcv package (Wood 2011) in R 4.1.2 (R Core Team 2022). The use of GAMs allows us to account for the non-linear effect of time of day on the soundscape by fitting a spline to the variable. All models included year and time of day as predictors, as well as a random effect of date. Year was included as a factor in the model, comparing sound during the 2020 zoo closure period to the period of zoo opening in 2019. Including a random effect of date accounts for non-independence of data points within days if days differ intrinsically. For analyses examining individual frequency bands the models also included the frequency band of the measurement (as a factor) and the interaction between frequency band and year. As different aviaries are expected to behave differently both within and between years, the set of models described above were run separately for each aviary in the dataset. This aids interpretation of the results by removing high order interactions between factors.

### 2.5 Data availability statement

Data and code associated with this manuscript can be found at https://doi.org/10.48420/22921460

## 3. Results

### 3.1 Broad frequency band measures

Broad frequency band measures of sound pressure levels varied significantly between years, although effects differed among aviaries. When examining dB(A), Sound pressure levels (SPLs) as measured in dB(A) were lower in the Dragons in Danger and Sumatra aviaries in 2020, in the absence of visitors, compared to 2019 (Table 1; Figure 1A). There was no significant difference in dB(A) in the Bali Temple aviary between years (Table 1; Figure 1A). Sound pressure levels measured as dB(Z) were significantly lower in all 3 aviaries in 2020 compared to 2019 (Table 1; Figure 1B). Across all models, the spline for time was significant (p<0.01), meaning that sound pressure levels varied significantly over the course of the day.

**Figure 1:**
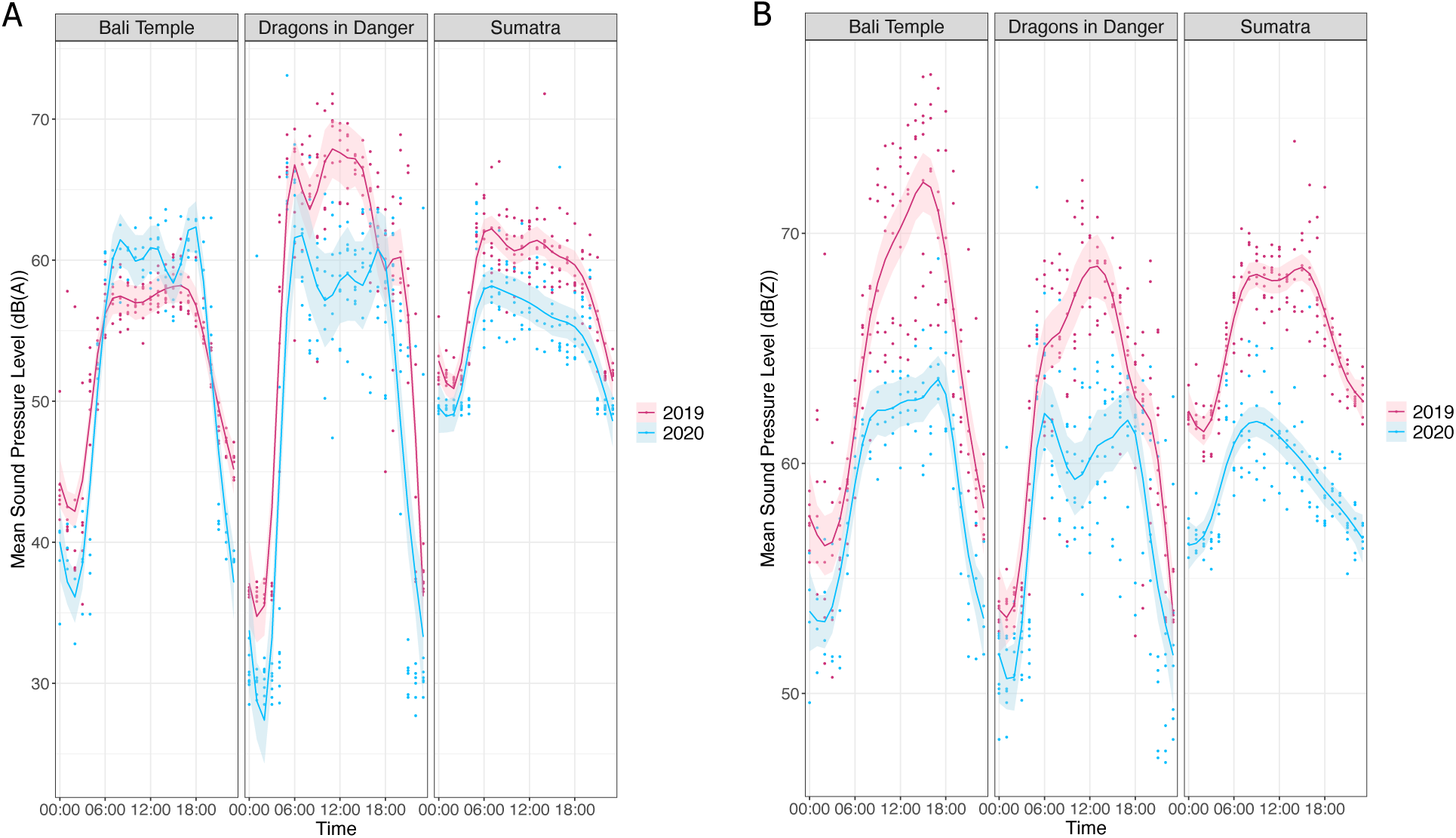
Daily patterns of sound pressure levels (broad frequency band) for zoo closure (2020, blue) and zoo open (2019, red) periods across three zoo aviaries. Panel A shows the mean sound pressure levels in dB(A). Panel B shows mean sound pressure levels in dB(Z). Lines show the predicted values (using GAMs) and the shaded area shows the standard error around the prediction. Individual points show raw data values.

**Table 1:**
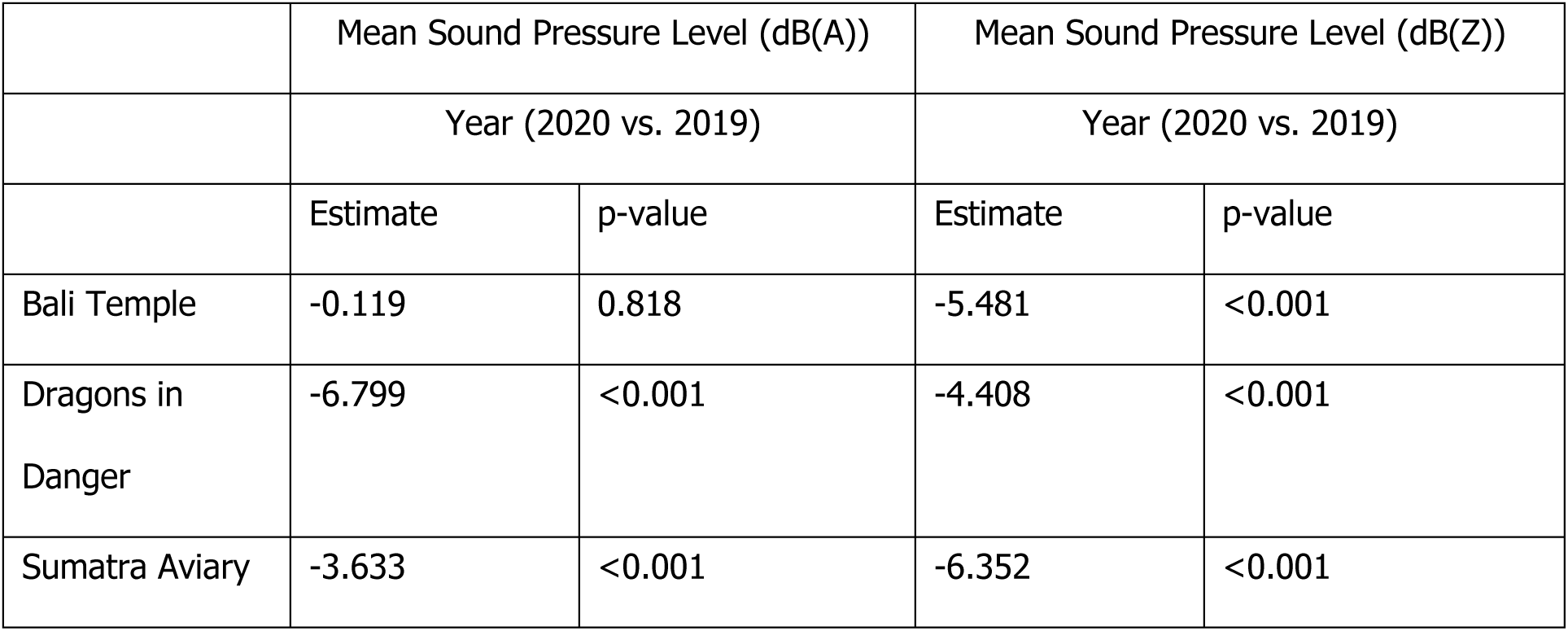
Generalized Additive Models (GAMs) examining the effect of year (as a factor) on mean sound pressure level (SPL) in dB(A) and dB(Z) in three Chester Zoo aviaries. Estimates represent the difference in 2020 compared to 2019

### 3.2 Defined frequency band measures

Changes in SPLs within defined bands varied among aviaries and frequency bands. As the models contained significant interactions, we present pairwise differences for frequency band and year to show how sound within each band changed between the zoo open and zoo closed periods. In all three aviaries, SPLs in the low and very low frequency bands decreased in 2020 compared to 2019 (Table 2; Figure 2). However, SPLs in the mid frequency band decreased only in the Sumatra and Dragons in Danger aviaries. SPLS in the high frequency band decreased in the Sumatra and Dragons in Danger aviaries, but increased in the Bali Temple aviary (Table 2; Figure 2). Across all models, the spline for time was significant (p<0.001), meaning that SPLs varied significantly over the course of the day.

**Figure 2:**
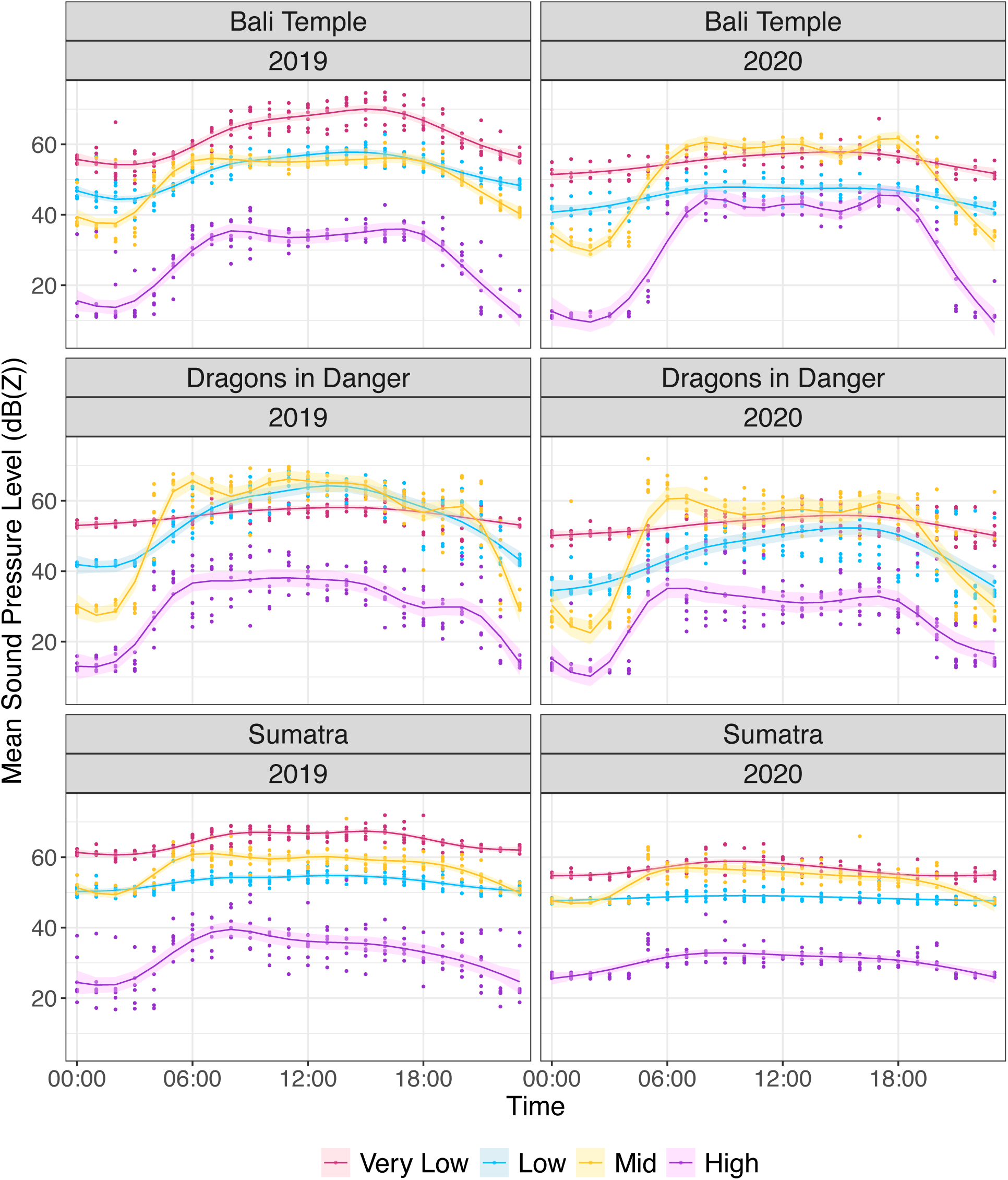
Mean sound pressure levels (dB(Z)) within frequency bands (Very Low (17.6-111.4 Hz), Low (111.4 – 890.9 Hz), Mid (890-9 – 8979.7 Hz), High (8979.7 – 11313.7 Hz)) for zoo closure (2020) and zoo open (2019) periods in three zoo aviaries. Lines show the predicted values and the shaded area (using GAMs) shows the standard error around the prediction. Individual points show raw data values.

**Table 2:**
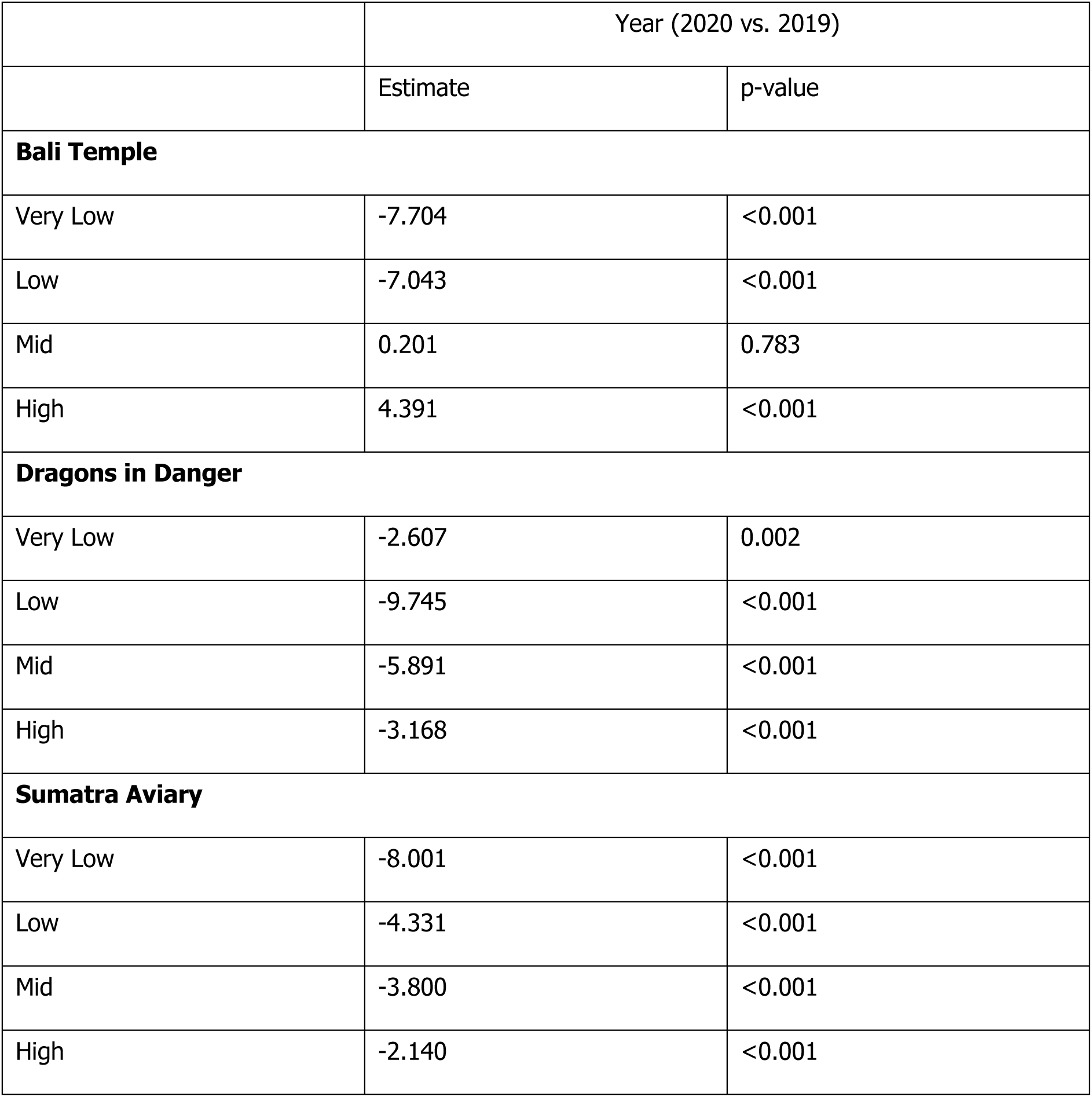
Generalized Additive Models (GAMs) examining the effect of year and frequency band (Very Low (17.6-111.4 Hz), Low (111.4 – 890.9 Hz), Mid (890-9 – 8979.7 Hz), High (8979.7 – 11313.7 Hz)) on sound pressure levels (dB(Z)) in three Chester Zoo aviaries. Values represent pairwise differences illustrating the change in the mean sound pressure level (SPL) within each frequency band during zoo closure in 2020 compared to zoo opening in 2019

### 3.3 Acoustic Indices

There was a significant effect of zoo closure on the Acoustic Complexity Index (ACI), although the direction of this effect differed among aviaries (Table 3; Figure 3A). ACI values were higher in 2020 than 2019 in the Bali Temple aviary, indicating a more complex soundscape. However, in the Dragons in Danger and Sumatra aviaries, ACI values were lower in 2020 than 2019, indicating reduced soundscape complexity in 2020. Across all models, the spline for time was significant (p<0.001), meaning that ACI values varied significantly over the course of the day.

**Figure 3:**
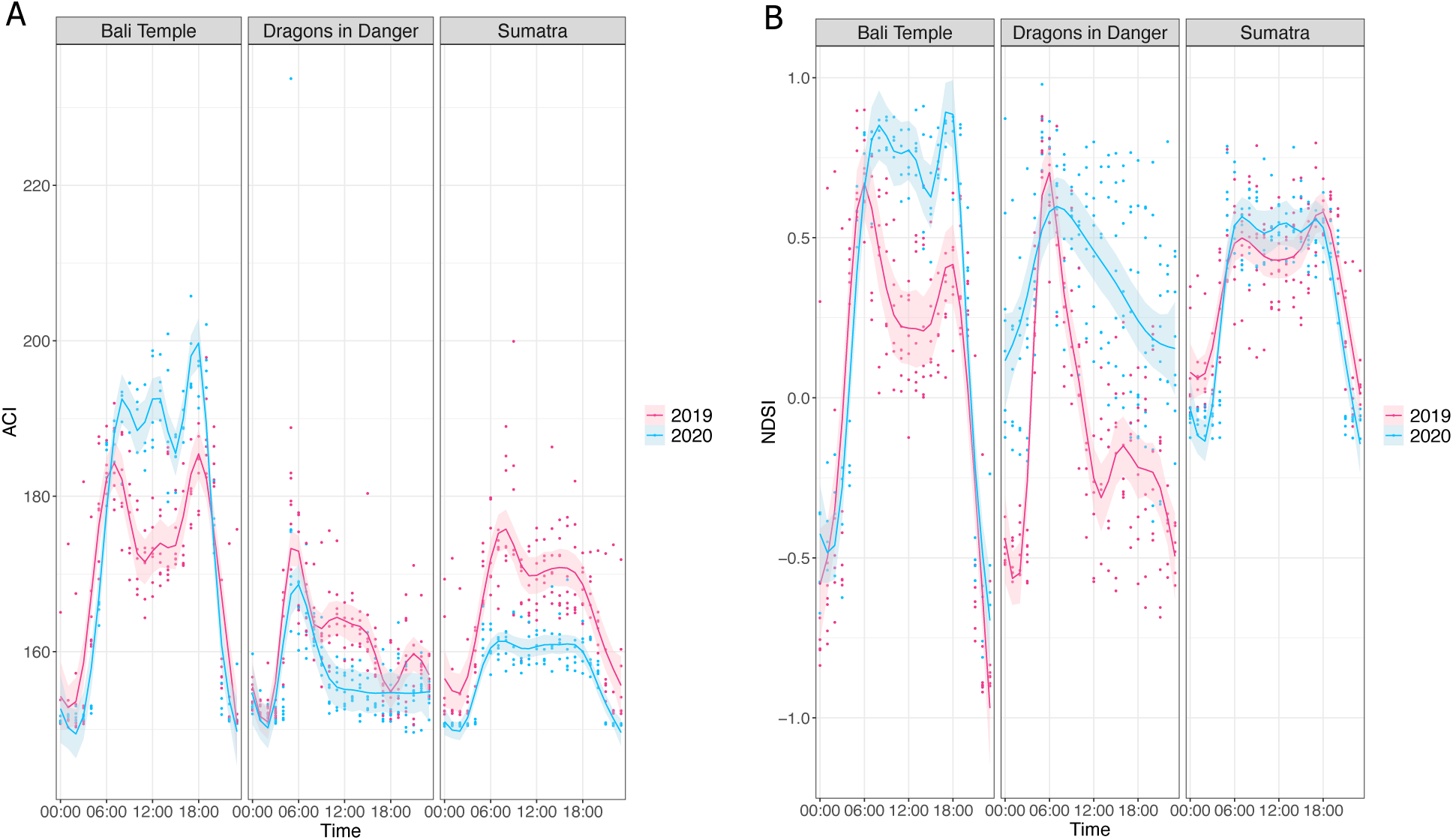
Daily patterns of acoustic indices in zoo aviaries for zoo closure (2020) and zoo open (2019) periods in three zoo aviaries. Panel A shows the Acoustic Complexity Index (ACI). Panel B shows the Normalized Difference Soundscape Index (NDSI). Lines show the predicted values and the shaded area shows the standard error around the prediction. Individual points show raw data values.

**Table 3:** Generalized Additive Models (GAMs) examining the effect of year (as a factor) on Acoustic Complexity Index (ACI) and Normalized Difference Soundscape Index (NDSI) in three zoo aviaries Estimates represent the difference in 2020 compared to 2019.

|  | ACI |  | NDSI |  |
| --- | --- | --- | --- | --- |
|  | Year (2020 vs. 2019) |  | Year (2020 vs. 2019) |  |
|  | Estimate | p-value | Estimate | p-value |
| Bali Temple | 5.549 | <0.001 | 0.252 | <0.001 |
| Dragons in Danger | -3.478 | 0.006 | 0.459 | <0.001 |
| Sumatra Aviary | -8.475 | <0.001 | -0.032 | 0.317 |

When considering the Normalized Difference Soundscape Index (NDSI), there was a significant effect of year in two out of the three aviaries examined. In the Bali Temple and Dragons in Danger aviaries, NDSI values were higher in 2020 than 2019 (Table 3; Figure 3B). This indicates a greater proportion of biotic compared to anthropogenic sound in the soundscape. However, there was no significant difference in NDSI values between years in the Sumatra aviary. Across all models, the spline for time was significant (p<0.001).

## 4. Discussion

We found significant differences in the soundscape of zoo aviaries between the zoo closure period in 2020 and a similar period of normal zoo operation in 2019. During the closure period, no visitors were present on the zoo site, and human activity surrounding the zoo was reduced due to the COVID-19 lockdown. Given the substantial reductions in human activity during 2020, it is apparent that chronic human presence in and around the zoo has a significant effect on the zoo soundscape, with aviaries being quieter, with less low frequency sound, and greater soundscape naturalness during the closure period.

Overall sound pressure levels (SPLs) across the whole frequency spectrum decreased during the zoo closure period. This pattern is similar to that reported by Quadros et al., (2014), who found that sound levels were higher on days when the zoo was open compared to when the zoo was closed across enclosures measured. Results were similar, but not identical, when considering A-weighted (dB(A)) and Z-weighted (dB(Z)) metrics; all three aviaries were significantly quieter in 2020 when considering dB(Z), but only two of the three aviaries showed a significant decrease in dB(A). These differences can be understood in more detail using the narrower frequency bands examined in our study. Low and very low frequency sound, which is down-weighted when calculating dB(A) to reflect human hearing sensitivity (Kurra 2021), was a significant component of the soundscape in all aviaries, and decreased significantly during the zoo closure period. Mid and high frequency sounds, which are given a larger weighting when calculating dB(A) (Kurra 2021), also decreased in two of the aviaries. However, mid frequency sound showed no difference and high frequency sound increased in the third aviary, which may contribute the differences between aviaries when using dB(A). Our results suggest that an unweighted metric, such as dB(Z), may be more useful when characterizing the effect of disturbances on the zoo soundscape, especially as weighted metrics, such as dB(A) may be inappropriate for assessing noise perceived by species with different hearing ranges to humans (Pater et al. 2009).

The use of defined, narrow frequency bands in addition to traditional, broad-band measures of SPLs can provide a more detailed overview of where sound energy is concentrated in the environment. This can help us to further explore information provided by other metrics, such as dB(A) and dB(Z), as well as identifying which species may be impacted most by disturbances. Species with sensitive hearing ranges or vocal ranges in frequency bands impacted by human presence may be disproportionately affected. The frequencies of sound examined (17.6 – 11313.17 Hz) in this study were appropriate for birds (Dooling 1992; Dooling et al. 2000; Catchpole & Slater 2008). However, many animals are sensitive to frequencies outside of this range, including infrasound (e.g., elephants (Payne et al. 1986)) and ultrasound (e.g., rodents (Sales 2010)), and defined bands could be tailored as necessary. Across aviaries, the loudest frequency bands differed, although low and very low frequency noise were prominent across aviaries during both years. Noise in these bands significantly decreased in 2020, indicating a significant effect of human presence. Birds vocalizing in this range, such as doves and parrots, may be particularly susceptible to human disturbances, with noise potentially masking important vocal signals or contributing to vocal change. Given the potential effects of noise on behaviour, welfare and conservation, zoos may wish to use mitigation measures to reduce the impact of sound on animals. Different materials may have different levels of effectiveness that vary between frequency bands (Orban et al. 2017). Therefore, measurements of SPLs in narrower frequency bands may be more beneficial for targeting mitigations and determining the effectiveness of such interventions. Our results suggest that mitigations targeting frequencies below 1000 Hz may be beneficial for reducing the impact of human disturbances on the soundscape.

We also examined the use of acoustic indices in zoo environments. Although acoustic indices can provide information about the characteristics of sound in the environment beyond its intensity (Sueur et al. 2014), they are not generally considered in captive environments (Clark & Dunn 2022). The Acoustic Complexity Index (ACI) is a measure of soundscape complexity, and correlates with animal vocal activity in wild environments (Pieretti et al. 2011). We did not find consistent differences in patterns of soundscape complexity between the two periods, with complexity increasing in one aviary and decreasing in the other two. This suggests that soundscape complexity, and the ACI, may not be a suitable measure to assess anthropogenic disturbances and the response of zoo-housed birds. Although not suitable as a measure of disturbance, ACI showed similar diel patterns to SPLs, increasing in the morning as more animals become active and decreasing towards the end of the day, indicating that peaks in ACI may still relate to vocal activity. In addition, the direction of change for ACI reflected changes in SPLs in the mid and high frequency bands. Where ACI values decreased, SPLs in both mid and high bands also decreased, and when the ACI value increased there was a corresponding increase in SPLs in the high frequency band. Most biotic sound is concentrated between 2 and 8 kHz (Pijanowski et al. 2011; Kasten et al. 2012), although some birds use higher frequencies in the vocal repertoire, so these changes in SPLs, and associated changes in ACI may indicate differences in biotic noise and vocal activity. Although vocal activity can shift due to anthropogenic noise disturbance (Brumm & Slabbekoorn 2005; Slabbekoorn & Ripmeester 2008; Ortega 2012), a number of other factors could have influenced our results. Vocal activity is likely to be affected by the time of year (Amrhein et al. 2002; Koloff & Mennill 2013; Digby et al. 2014), weather (O’Connor & Hicks 1980; Digby et al. 2014; Vokurková et al. 2018), and number of birds in and around the aviary. Furthermore, in complex and frequently disturbed captive environments, soundscape complexity may be influenced by additional factors beyond birds’ vocal activity, in particular the presence of human voices. Whilst soundscape complexity and the ACI could be useful for detecting peaks in vocal activity in birds, e.g., during breeding, to inform management decisions, further investigation is required to fully understand possible factors that influence the ACI in zoos and the relevance of soundscape complexity in captive environments.

The Normalized Difference Soundscape Index (NDSI) is a measure of soundscape ‘naturalness’ (Kasten et al. 2012). NDSI values were higher in 2020 for two of the three aviaries, indicating a more natural soundscape, with a greater proportion of ‘biotic’ (2-8 kHz) compared to ‘anthropogenic’ (1-2 kHz) noise. Given the decrease in human activity in 2020, the NDSI appears to be a successful indicator of disturbance in zoos, and could be used to determine the relative impact of human-generated sound. Whilst NDSI appears to be potentially useful in zoos, there are some potential limitations. In particular, our analyses of sound pressure levels in defined bands indicate that sounds below 1000 Hz are a significant component of the zoo soundscape, and these sounds decreased during the zoo closure period. However, when using the default settings, these frequencies are not used in the calculation of the NDSI, and so are not captured by the index, which may limit our understanding of overall disturbance. Despite this, NDSI values suggested a similar, but not identical, pattern of human disturbance to the soundscape as the SPLs in two of the three aviaries, but did not differ between years in the third. However, the NDSI provides information about relative changes in the soundscape, rather than changes in absolute values. So, if noise in both the ‘biotic’ and ‘anthropogenic’ bands change in the same way relative to one another, NDSI values will not change despite an overall change in SPLs. One of the most important limitations of the NDSI in zoos is the potential impact of human voices, as human speech covers a broad range of frequencies (Fant 2004), spanning both the ‘anthropogenic’ and ‘biotic’ bands of the default NDSI settings. Whilst this does not preclude the use of NDSI, further investigation would be beneficial to understand the usefulness of NDSI in monitoring zoo sound.

Across all measures, aviaries varied in terms of the soundscape and changes therein. Such differences may relate to aviary position within the zoo, setting (indoor or outdoor), popularity, and species composition. Previous studies have reported that indoor environments differed significantly from those outdoors (Pelletier et al. 2020). However, we did not find consistent differences between the indoor and outdoor aviaries. Changes in species composition and number of birds in aviaries between years may have significantly impacted our measures. For example, in the Bali Temple aviary, high frequency noise and ACI values increased, and mid frequency noise remained unchanged, despite SPLs in these bands decreasing in the other aviaries. Compared to the other aviaries, the Bali Temple had the largest population increase (73 birds in 2019 vs. 83 birds in 2020), which may have impacted biotic noise and vocal activity. In order to identify specific features that could impact changes in the soundscape, a larger sample size of aviaries would be required. However, it is possible that enclosures are uniquely impacted by different factors, in which case, generalizations would not be possible and enclosures would need to be studied on a case-by-case basis.

## 5. Conclusion

Differences in multiple measures of sound between a closure period in 2020 compared to a period of normal operation in 2019 indicated that human presence significantly affects the soundscape in zoos. We demonstrated that acoustic indices could be calculated in captive settings, and may be relevant for assessing changes and informing management. In particular, the NDSI captured some useful information about human disturbances. Although the measures used in our study provide a similar picture of human disturbance on the soundscape, different measures showed different patterns within and between aviaries. The use of multiple features of the soundscape, as in this study, can provide additional information beyond the traditional measures of ‘loudness’ in zoos, including the frequencies where sound energy is concentrated, relative differences between frequency bands, and overall characteristics of the soundscape. Therefore, the use of multiple measures can provide a more comprehensive overview of the soundscape and allow for a more in-depth exploration into the effects of disturbances, and which species may be most affected and require intervention. Our study suggests that mitigation measures to reduce the impact of sound disturbance from humans could alter the soundscape experienced by birds and other taxa by reducing low frequency noise. Moving forwards, further examination of acoustic indices in captivity, such as how they are impacted by features such as human speech, would be beneficial to fully understand their value for describing and assessing the zoo soundscape.

## 6. Author Contributions

RL and LW conceptualized the study, with all authors contributing to refining data extraction protocols. RL and LW were involved in data collection for the project. RL completed the extraction of sound measures. RL and TG were responsible for data analysis. RL wrote the initial draft of the manuscript, including data visualizations, with all authors contributing to editing the manuscript. LW, SdK, and TG provided supervision to RL throughout the course of the project.

## Acknowledgements

RL’s work on this project was funded by the Natural Environment Research Council (NERC) EAO Doctoral Training Partnership (grant NE/L002469/1) supported by the Chester Zoo Conservation Scholar and Fellow Scheme. We would like to thank the bird teams at Chester Zoo for facilitating with the data collection for this project.

