## Supplementary Material 1 for "Assessing the effect of zoo closure on the soundscape using multiple measures"

Supplementary Material S1: Species lists for the three zoo aviaries in which the sound environment was investigated (Bali Temple, Dragons in Danger, Sumatra Aviary) in 2019 during zoo opening and 2020 during zoo closure. Date ranges of recordings are indicated in brackets.

**Bali Temple**

| ***2019 (30/4/2019 – 7/5/2019)*** | | | ***2020 (08/04/2020- 12/04/2020)*** | | |
|---|---|---|---|---|---|
| ***Species Name*** | ***Latin Name*** | ***Number*** | ***Species Name*** | ***Latin Name*** | ***Number*** |
| Bali myna | *Leucopsar rothschildi* | 9 | Bali myna | *Leucopsar rothschildi* | 5 |
| Java sparrow | *Lonchura oryzivora* | 55 | Java sparrow | *Lonchura oryzivora* | 65 |
| Pied imperial pigeon | *Ducula bicolor* | 5 | Magpie robin | *Copsychus saularis* | 2 |
| Purple-naped lory | *Lorius domicella* | 2 | Pied imperial pigeon | *Ducula bicolor* | 5 |
| Sumatran laughing thrush | *Garrulax bicolor* | 1 | Purple-naped lory | *Lorius domicella* | 4 |
| Yellow-backed chattering lory | *Loris garrulus flavopalliatus* | 1 | Sumatran laughing thrush | *Garrulax bicolor* | 1 |
| | | | Yellow-backed chattering lory | *Loris garrulus flavopalliatus* | 1 |

**Sumatra aviary**

| ***2019 (07/05/2019 – 14/5/2019)*** | | | ***2020 (22/04/2020 – 28/04/2020)*** | | |
|---|---|---|---|---|---|
| ***Species Name*** | ***Latin Name*** | ***Number*** | ***Species Name*** | ***Latin Name*** | ***Number*** |
| Asian glossy starling | *Aplonis panayensis* | 24 | Asian glossy starling | *Aplonis panayensis* | 26 |
| Bronze-tailed peacock pheasant | *Polyplectron chalcurum* | 2 | Bronze-tailed peacock pheasant | *Polyplectron chalcurum* | 2 |
| Chestnut-backed thrush | *Geokichla dohertyi* | 4 | Chestnut-backed thrush | *Geokichla dohertyi* | 5 |
| Chestnut-bellied tree partridge | *Arborophila javanica* | 1 | Chestnut-bellied tree partridge | *Arborophila javanica* | 1 |
| Emerald dove | *Chalcophaps indica* | 9 | Emerald dove | *Chalcophaps indica* | 6 |
| Fairy bluebird | *Irena puella* | 1 | Fairy bluebird | *Irena puella* | 1 |
| Fire-tufted barbet | *Psilopogon pyrolophus* | 1 | Fire-tufted barbet | *Psilopogon pyrolophus* | 1 |
| Javan green magpie | *Cissa thalassina* | 3 | Javan green magpie | *Cissa thalassina* | 2 |
| Magpie robin | *Copsychus saularis* | 1 | Salvadori's pheasant | *Lophura inomata* | 2 |
| Salvadori's pheasant | *Lophura inomata* | 2 | Silver-eared mesia | *Leiothrix argentauris* | 7 |
| Silver-eared mesia | *Leiothrix argentauris* | 3 | | | |

**Dragons in Danger**

| ***2019 (14/05/2019 – 21/05/2019)*** | | | ***2020 (30/4/2020 – 08/05/2020)*** | | |
|---|---|---|---|---|---|
| ***Species Name*** | ***Latin Name*** | ***Number*** | ***Species Name*** | ***Latin Name*** | ***Number*** |
| Black-naped fruit dove | *Ptiliopus melanospilus* | 3 | Black-naped fruit dove | *Ptiliopus melanospilus* | 4 |
| Cinnamon ground dove | *Gallicolumba rufigula* | 4 | Cinnamon ground dove | *Gallicolumba rufigula* | 3 |
| Fairy bluebird | *Irena puella* | 3 | Fairy bluebird | *Irena puella* | 2 |
| Great argus | *Argusianus argus* | 2 | Great argus | *Argusianus argus* | 2 |
| Luzon bleeding heart dove | *Gallicolumba luzonica* | 6 | Javan green magpie | *Cissa thalassina* | 2 |
| Malayan great argus | *Argusianus argus argus* | 1 | Luzon bleeding heart dove | *Gallicolumba luzonica* | 6 |
| Mindanao bleeding heart dove | *Gallicolumba criniger* | 2 | Malayan great argus | *Argusianus argus argus* | 1 |
| Montserrat oriole | *Icterus oberi* | 2 | Mindanao bleeding heart dove | *Gallicolumba criniger* | 3 |
| Palawan peacock pheasant | *Polyplectron superbus* | 2 | Palawan peacock pheasant | *Polyplectron superbus* | 1 |
| Philippine mouse-deer | *Tragulus nigricans* | 1 | Philippine mouse-deer | *Tragulus nigricans* | 1 |
| Pink-headed fruit dove | *Ptilinopus porphyrea* | 1 | Pink-headed fruit dove | *Ptilinopus porphyrea* | 1 |
| Sumatran laughing thrush | *Garrulax bicolor* | 2 | Superb fruit dove | *Ptilinopus superbus* | 5 |
| Superb fruit dove | *Ptilinopus superbus* | 4 | Victoria crowned pigeon | *Goura victoria* | 1 |
| Visayan tarictic hornbill | *Penelopides panini panini* | 5 | Visayan tarictic hornbill | *Penelopides panini panini* | 2 |
| White-naped pheasant pigeon | *Otidiphaps aruensis* | 2 | White-naped pheasant pigeon | *Otidiphaps aruensis* | 2 |
